# A hybrid demultiplexing strategy that improves performance and robustness of cell hashing

**DOI:** 10.1101/2023.04.02.535299

**Authors:** Lei Li, Jiayi Sun, Yanbin Fu, Siriruk Changrob, Joshua J.C. McGrath, Patrick C. Wilson

**Author notes:** Correspondence (P.C.W.). These authors contributed equally (Lei Li, Jiayi Sun, Yanbin Fu).

## Abstract

Recent advances in single cell RNA sequencing allow users to pool multiple samples and demultiplex in downstream analysis, which greatly increase experimental efficiency and cost-effectiveness. Among all the demultiplexing methods, nucleotide barcode-based cell hashing has gained widespread popularity due to its compatibility and simplicity. Despite these advantages, certain issues of this technic remain to be solved, such as challenges in distinguishing true positive from background, high reagent cost for samples with large cell numbers, and unpredictable false negative and false doublet rates. Here, we propose a hybrid demultiplexing strategy that increases calling accuracy and cell recovery of cell hashing without adding experimental cost. In this approach, we computationally cluster all single cells based on their natural genetic variations and assign donor identity by finding the dominant hashtag in each genotype cluster. This hybrid strategy assigns donor identity to any cell that is identified as singlet by either genotype clustering or cell hashing, which allows us to demultiplex most majority of cells even if only a small fraction of cells are labeled with hashtags. When comparing its performance with cell hashing on multiple real-world datasets, this hybrid approach consistently generates reliable demultiplexing results with increased cell recovery and accuracy.

**Key Points:**

1. The improved cut-off calling tool, HTOreader, accurately distinguishes true positive from background signal for each individual hashtag.
2. The hybrid demultiplexing strategy increases cell recovery of cell hashing by increasing cut-off calling accuracy and decreasing false negative and false double rates.
3. The hybrid strategy enhances cost-effectiveness of cell hashing and consistently produces reliable demultiplexing results, regardless of hashtag staining quality.
4. The hybrid strategy can be seamlessly integrated into a variety of single-cell experimental protocols and analytic pipelines.

## Introduction

The technical advances in single-cell sequencing have greatly benefited biological and medical research by enhancing our ability to investigate cellular mechanisms of homeostasis and disease in a more precise, high-resolution, and multi-omic fashion(Tang et al., 2009, Svensson et al., 2018, Stuart and Satija, 2019). In the past decade, more and more single-cell methods have been proposed to improve the quality, magnitude, modality, and economy of single-cell experimental approaches(Tang et al., 2009, Picelli et al., 2013, Klein et al., 2015, Macosko et al., 2015, Cao et al., 2017, Stoeckius et al., 2018, Kang et al., 2018, Heaton et al., 2020). Among them, single cell sample pooling and demultiplexing can greatly reduce the per-cell cost, and therefore have been extensively studied.

Sample-multiplexing approaches in single-cell sequencing have been proposed firstly in 2015 and grown explosively (Zhang et al., 2022). To date, as summarized by a previous publication, there are at least 31 sample-multiplexing approaches in single-cell biology study (Zhang et al., 2022). For single-cell RNA-seq, existing methods fall into several major technical solutions, including nucleotide barcode-based methods, natural genetic variation-based methods, and vector-based barcoding methods. Among them, nucleotide barcode-based methods, such as cell hashing (utilize oligo-tagged antibodies), MULTI-seq (utilize lipid-tagged indices), ConA-based sample-barcoding strategy (CASB) and ClickTags (Stoeckius et al., 2018, McGinnis et al., 2019, Fang et al., 2021, Gehring et al., 2020), as well as natural genetic variation-based methods, such as demuxlet, scSplit, Vireo, and Souporcell (Xu et al., 2019, Huang et al., 2019, Kang et al., 2018, Heaton et al., 2020) have been widely used in a variety of single-cell experiments. Mechanistically, nucleotide barcode-based methods link sequences of unique oligonucleotide tags to sample identity, and those oligonucleotides can be either anchored to cellular or nuclear membranes (e.g. Cell hashing), internalized to the cytoplasm or nucleus (e.g. SBO), or incorporated during library construction (e.g. BART-Seq)(Stoeckius et al., 2018, Shin et al., 2019, Uzbas et al., 2019). On the other hand, natural genetic variation-based methods identify single nucleotide polymorphisms (SNPs) of each donor from sequencing data to determine donor identity.

Based on Cellular Indexing of Transcriptomes and Epitopes by Sequencing (CITE-seq) technology, cell hashing has been one of the most commonly used demultiplexing approaches due to its compatibility and simplicity. It involves staining cells from each sample with uniquely barcoded antibodies [also called hashtags, or hashtag oligonucleotides (HTOs)] that recognize ubiquitously expressed surface antigens, such as β2-microglobulin (Stoeckius et al., 2017). The labeled cells can then be pooled and prepared for single cell sequencing, and sample identities of single cells can be determined based on relative enrichment level of each hashtag using computational approaches. Despite the widespread usage, cell hashing has certain limitations that need to be overcome. Firstly, it is challenging to accurately distinguish true positive from background signal for each individual hashtag. Secondly, the maximum recovery rate of cell hashing is limited by high false doublets rate. Even for a benchmark dataset where hashtag staining quality is high, when using eight hashtags, only about 80% of cells are identified as singlets, which is lower than 90% recovery generated by genetic variation-based methods, such as Souporcell (Stoeckius et al., 2018; Heaton et al., 2020). Thirdly, the performance of cell hashing is highly sensitive to the staining quality of the hashtags. Instances of hashtag failure occur occasionally, leading to a high percentage of false negative cells (Dugan et al., 2021). Lastly, cell hashing can be expensive when processing samples with large number of cells.

Here, we propose a hybrid single-cell demultiplexing strategy that improves overall performance by combining cell hashing with genetic variation-based methods. We set out to improve the existing cut-off calling algorithm for cell hashing by developing HTOreader, a tool that accurately distinguish true positive signals from background for each individual hashtag. By integrating results of HTOreader and Souporcell on multiple datasets, we observe an increase in cell recovery rate for all datasets when compared to original cell hashing approach, and almost all datasets achieve 90% singlet rate. As expected, the pronounced increases occur in datasets that have low recovery due to poor hashtag staining quality, indicating performance of this hybrid approach is not subject to staining quality. Since the generation of genotyping clusters is independent of cell hashing data, we apply our strategy to multiple integrated datasets with cells from the same group of donors that do or do not have hashtag data. The results demonstrate that the performance remain consistent even if a large fraction of cells are not stained with hashtags, indicating reagent costs can be substantially decreased by only staining a small amount of cells from each donor. Finally, by integrating results from methods using completely different mechanisms, the two methods can automatically validate each other so that potential errors arise from any of them can be easily identified, making the hybrid strategy more reliable than single-modal methods. Together, our hybrid strategy provide a robust, accurate, and cost-effective solution for demultiplexing single-cell RNA sequencing results.

## Results

### Limitations of cell hashing in performance and economy

There are several factors that influence the performance of cell hashing, including cut-off calling accuracy, false doublet rate, and false negative rate. Our analysis on multiple real-world datasets reveal varying singlet rates: 81.67% (dataset Stoeckius-2018, Figure 1-A), 79.57% (dataset 3V007, Figure 1-B), 80.31% (dataset S414, Figure 1-C), 62.29% (dataset R125, Figure 1-D), and 30.56% (dataset R6, Figure 1-E). Firstly, even in the benchmark dataset with high-quality staining and a well-balanced distribution of cells in each sample (dataset Stoeckius-2018, Figure 1-A), we observed 16.33% of cells were identified as doublets, which include real doublets that consist of multiple individually-hashtagged cells sticking together, and false doublets that consist of single cells labeled with multiple hashtags. The inability to distinguish real doublets (which must be excluded from downstream analysis) from false doublets (which should be included if they pass quality control) results in substantial cell loss. Furthermore, 13.26%, 7.51%, and 31.04% of cells in datasets 3V007, S414, and R125 were identified as negative. Surprisingly, a large portion of these negative cells were singlets with low hashtag signal. We found that the built-in algorithm in Seurat is unable to accurately distinguish positive signal from background, and therefore incorrectly categorize these cells as negative (“negative” cells in Fig 1-B,C, and D have sufficient signal to hashtag1, hashtag3, and hashtag2, respectively). Finally, 68.98% of cells in dataset R6 were identified as negative due to poor staining quality of hashtags 3 and 4, rendering the longitudinal information for most cells in this dataset inaccessible (Figure 1-E). Under this situation, hashtag identities of cells that are not properly stained and only have background level of signal are lost unavoidably.

**Figure 1.**
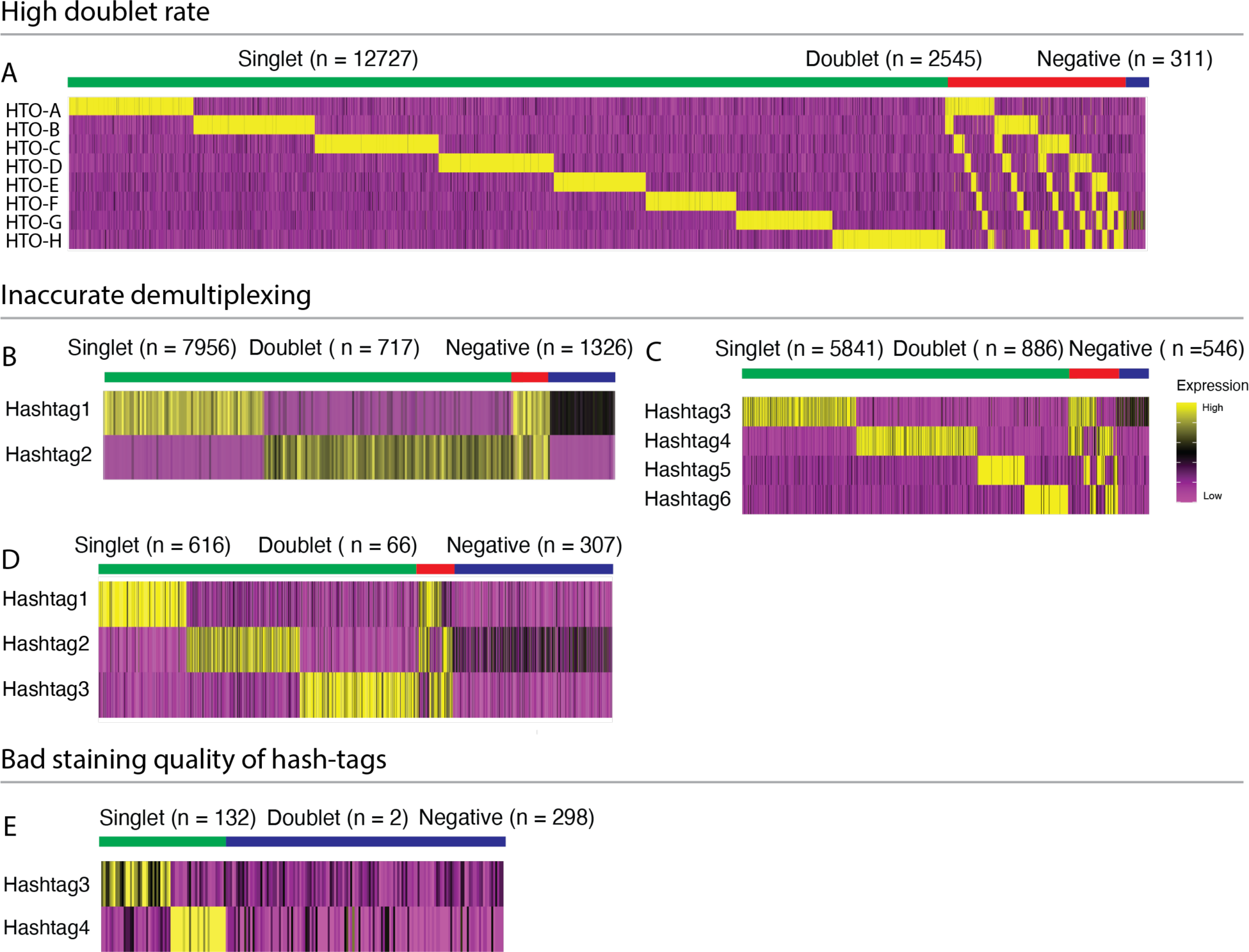
Limitations of cell hashing method revealed by real-world datasets. Expression of hashtags in datasets Stoeckius-2018, 3V007, S414, R125 and R6 are shown by heatmaps. Singlet, doublet, and negative groups were indicated by a color bar with number of cells in each group. Panels A – E share the same scale bar of heatmap to the right. Five datasets were grouped by three reasons of cell loss in cell hashing: high doublet rate, inaccurate demultiplexing due to insufficient hashtag signals, and bad staining quality of one or more hashtags. (**A**) Demultiplexing of dataset Stoeckius-2018. (**B**) Demultiplexing of dataset 3V007. (**C**) Demultiplexing of dataset S414. (**D**) Demultiplexing of dataset R125. (**E**) Demultiplexing of dataset R6.

In addition to performance, experimental cost is another factor that limits the applicability of cell hashing. In the current protocol, all cells must be stained with a hashtag antibody labeled with a unique barcode that correlate with their sample identity. Consequently, the reagent cost is directly tied to the total number of cells that need to be stained, which can be expensive for massive datasets and large-scale studies. In conclusion, efforts that improve the cell recovery, accuracy, robustness, and cost-effectiveness of cell hashing is desperately needed for advancement of single-cell analysis.

### HTOreader, an improved cut-off calling method that increases accuracy of cell hashing

One of the major challenges in cell hashing is to accurately determine cutoffs that distinguish true positive signals from background for each individual hashtag. To address it, we developed a finite-mixture-modeling-based pipeline, HTOreader, aiming to enhance cutoff calling accuracy. The pipeline involves four key steps: 1) Normalizing raw counts for each individual hashtag, 2) Employing a mixture model to fit the normalized data into two Gaussian distributions, representing background and true positive groups, 3) Determining the optimal cutoff based on the means and standard deviations of the two groups, 4) Assigning sample identities to each individual cell (singlet, doublet, or negative) by assessing their hashtag profile (Figure 2-A). To make this tool user-friendly, we have ensured its compatibility with Seurat, enabling seamless integration into popular existing pipelines.

**Figure 2.**
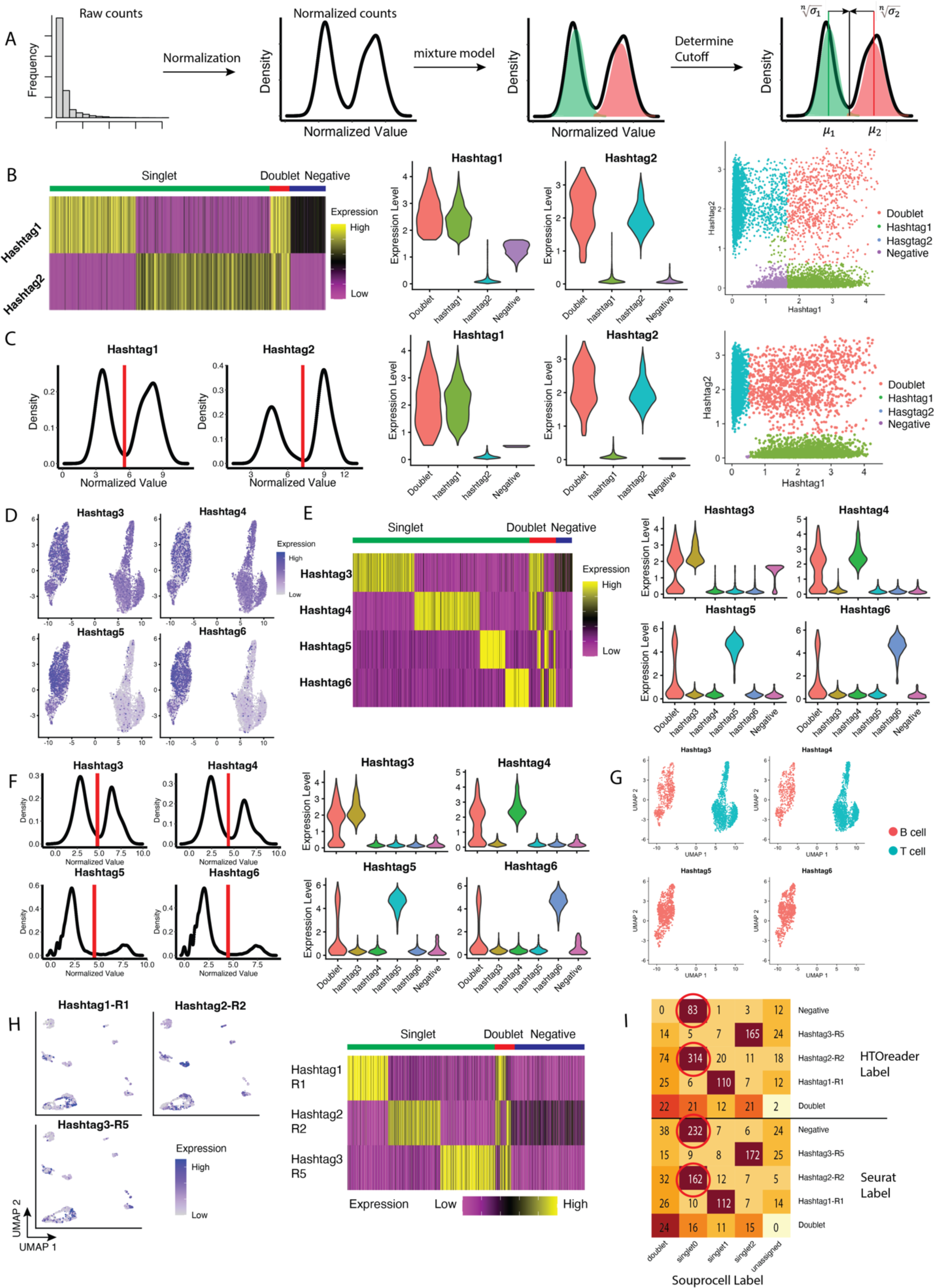
HTOreader achieves high accuracy and increases cell recovery for cell hashing datasets. (**A**) The workflow of HTOreader to determine proper cutoff for each cell hashtag. (**B**) Cell demultiplexing by Seurat on 3V007. Left: expression heatmap of both hashtags with Seurat labels annotation. Center: violin plot of expression of both hashtags on Seurat groups. Right: Scatter plot of hashtag expression with Seurat label color coding. (**C**) Cell demultiplexing by HTOreader on dataset2. Left: cutoffs determined by HTOreader based on density of normalized values for both hashtags; Center: violin plot of expression of both hashtags on HTOreader groups; Right: Scatter plot of hashtag expression with HTOreader label color coding. (**D**) Expression of four hashtags on the UMAP embedding of S414. (**E**) Cell demultiplexing by Seurat on S414. Left: expression heatmap of all hashtags with Seurat labels annotation. Right: violin plot of expression of all hashtags on Seurat groups. (**F**) Cell demultiplexing by HTOreader on S414. Left: cutoffs determined by HTOreader based on density of normalized values for all hashtags; Right: violin plot of expression of all hashtags on HTOreader groups. (**G**) Cells labeled by different hashtags demultiplexed by HTOreader. (**H**) Cell demultiplexing by Seurat on R125. Left: Expression of three hashtags on the UMAP embedding of dataset4. Right: expression heatmap of all hashtags with Seurat labels annotation. (**I**) Comparing HTOreader and Seurat using Souprocell as benchmark on R125. Numbers of cells are indicated in the heatmap.

To evaluate the performance of HTOreader, we set out to run a perfectly balanced benchmark dataset (Stoeckius-2018). Results demonstrated that the performance of HTOreader was comparable to the demultiplexing function in Seurat (Figure S1). Next, we compared the performances of these two methods on two unpublished datasets (3V007, S414) that contain B and T cell data with imbalanced sample sizes (Figure 2-B,E). For 3V007, Seurat alone failed to determine a proper cutoff for hashtag1 and incorrectly assigned 1322 hashtag1-labeled cells into the negative group, resulting in a 13.22% cell loss (Figure 2-B). In contrast, by generating accurate cutoffs with HTOreader, the singlet rate increased from 79.57% to 88.71% (Figure 2-C). For S414, as we expected, Seurat incorrectly assigned 435 hashtag3-labeled cells into negative group, resulting in a 5.98% cell loss (Figure 2-D,E). Again, HTOreader was able to increase the singlet rate from 80.31% to 84.74% (Figure 2-F,G). To further demonstrate that this issue commonly exists, we performed the comparison on a published dataset (R125, Figure 2-H,I)(Dugan et al., 2021). Unsurprisingly, Seurat mislabeled 210 hashtag2_R2-labeled cells (21.23% of total cell) and HTOreader increased the singlet rate from 62.29% to 82.10% (Figure 2-H). To verify the accuracy of results generated by HTOreader, we ran R125 via Souporcell, a highly accurate, genotype-based demultiplexing pipeline (Heaton et al., 2020). Results showed that almost all hashtag2_R2-labeled cells retrieved from negative group by HTOreader were labeled as “singlet0” in Souporcell, demonstrating the superior accuracy of HTOreader (Figure 5-I). Therefore, compared to Seurat, HTOreader has similar performance for well-balanced datasets and much higher accuracy and cell recovery rate when running imbalanced, real-world datasets.

Next, we ran multiple real-world datasets (8pool-CA, S414, 3V007, and R125) that HTOreader successfully demultiplexed via several existing cell hashing-based methods, including MULTI-seq, HTOdemux, GMM-Demux and two BFF models (McGinnis et al., 2019, Hao et al., 2021, Xin et al., 2020, Boggy et al., 2022). Results on 8pool-CA showed that the two BFF models (BFF_raw and BFF_cluster) failed to identify cells from sample S282 (Table S1). The GMM-Demux method incorrectly clustered more than half of the cells into “doublet” clusters and only identified 39.82% of cells as singlet. Results on dataset S414 suggested that all methods achieved satisfactory performance (Table S2). Results on dataset 3V007 demonstrated that despite all methods achieved high cell recovery rate, HTOdemux failed to retrieve hashtag1-labeled cells from negative group (Table S3). For dataset R125, MULTI_seq produced low singlet rate (45.50%) because almost 1/3 of total cells were incorrectly categorized as negative (Table S4). To conclude, HTOreader consistently demonstrates robust and accurate performance across multiple real-world datasets, whereas other methods generate inaccurate results for at least one of the datasets (Table 1).

**Table 1.**
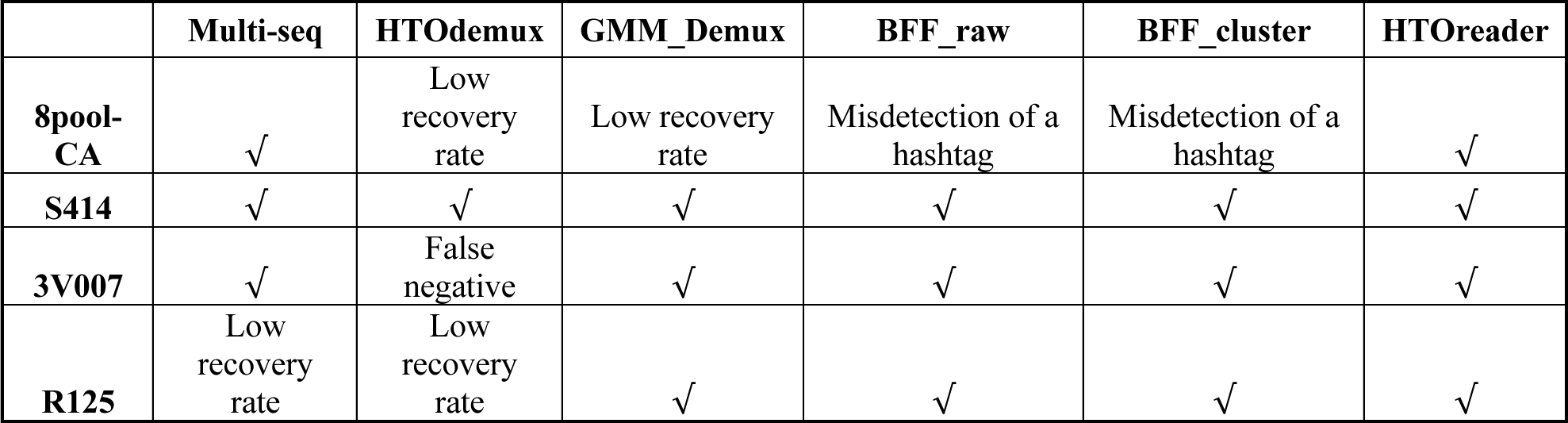
Comparisons of existing cell hashing demultiplexing methods on multiple real-world datasets.

Lastly, it is worth noting that many existing cell hashing-based methods, such as GMM-Demux, are written in Python and therefore are not directly accessible for the R community. Recently, Boggy et al. introduced BFF model and developed cellhashR, an R package that implements several popular methods, which comes in handy for R community (Boggy et al., 2022). However, among all these methods, only HTOreader and HTOdemux (method used by Seurat) offer direct compatibility with Seurat pipeline. Thus, HTOreader stands out as a convenient, accurate, and robust cell hashing-based demultiplexing tool catered specifically for R users, especially for those who utilize Seurat for their single-cell analysis.

### A hybrid demultiplexing strategy that increases cell recovery and cost-effectiveness of cell hashing

Although HTOreader improves performance of cell hashing by drawing more accurate cut-offs, cell loss caused by poor staining quality and high false-doublet rate remains unfixed. To address these issues, we exploit the recent advances in genetic variation-based demultiplexing methods, which clusters single cells based on their SNPs. Here, we propose a hybrid demultiplexing strategy that integrates cell hashing and SNP profiles of each individual cell. Using this hybrid strategy, sample identities of single cells that are poorly stained (false negative) or stained by multiple hashtags (false doublet) can be determined by the hashtag of those singlets that they cluster with based on SNPs. Notably, this hybrid strategy is compatible with all existing cell hashing-based (e.g. GMM-Demux, demuxEM, Seurat HTODemux, and BFF) and genetic variation-based (e.g. scSplit, Vireo, and Souporcell) demultiplexing methods. In the following analysis, we use HTOreader and Souporcell as a representative combination.

To demonstrate its performance, we applied this hybrid strategy to an unpublished real-world dataset (8pool). In this dataset, a small aliquots of PBMCs from each of eight donors were stained with hashtag antibodies individually and pooled together to sort carrier CD19+ B cells and CD4+ T cells (8pool-CA), whereas the rest of PBMCs were pooled together to sort antigen-specific B cells (8pool-AS) (Figure S2-A,B). We ran all cells in 8pool-CA and 8pool-AS through Souporcell, and cells in 8pool-CA through HTOreader. Results showed that all hashtags were stained with high quality and HTOreader divided all cells in 8pool-CA into ten groups, including 10,000 cells in eight singlet groups (77.6%), 2,759 cells in doublet (21.41%) and 127 cells in negative (0.98%) (Figure 3-A, Figure S2-C,D, Table 2). On the other hand, Souporcell divided cells in 8pool-CA into ten genotype clusters, 11,552 cells in eight singlet clusters (89.65%), 1,326 cells in doublet cluster (10.29%) and 8 cells in unassigned cluster (0.06%). As expected, the singlet rate of 8pool-CA by Souporcell reached 89.65%, which is 12.05% higher than that of cell hashing (Figure 3-B, Table 2). To take a closer look, 98.88% of singlets identified by HTOreader were also registered as singlet in Souporcell, validating the accuracy of both methods (Figure 3-B). Within the extra 12.05% singlets recovered by Souporcell, a small fraction were identified as negative whereas most majority were identified as doublet by HTOreader (1,541 out of 1664), consistent with the observation of high false-doublet rate in cell hashing (Figure 3-B, Table 2).

**Figure 3.**
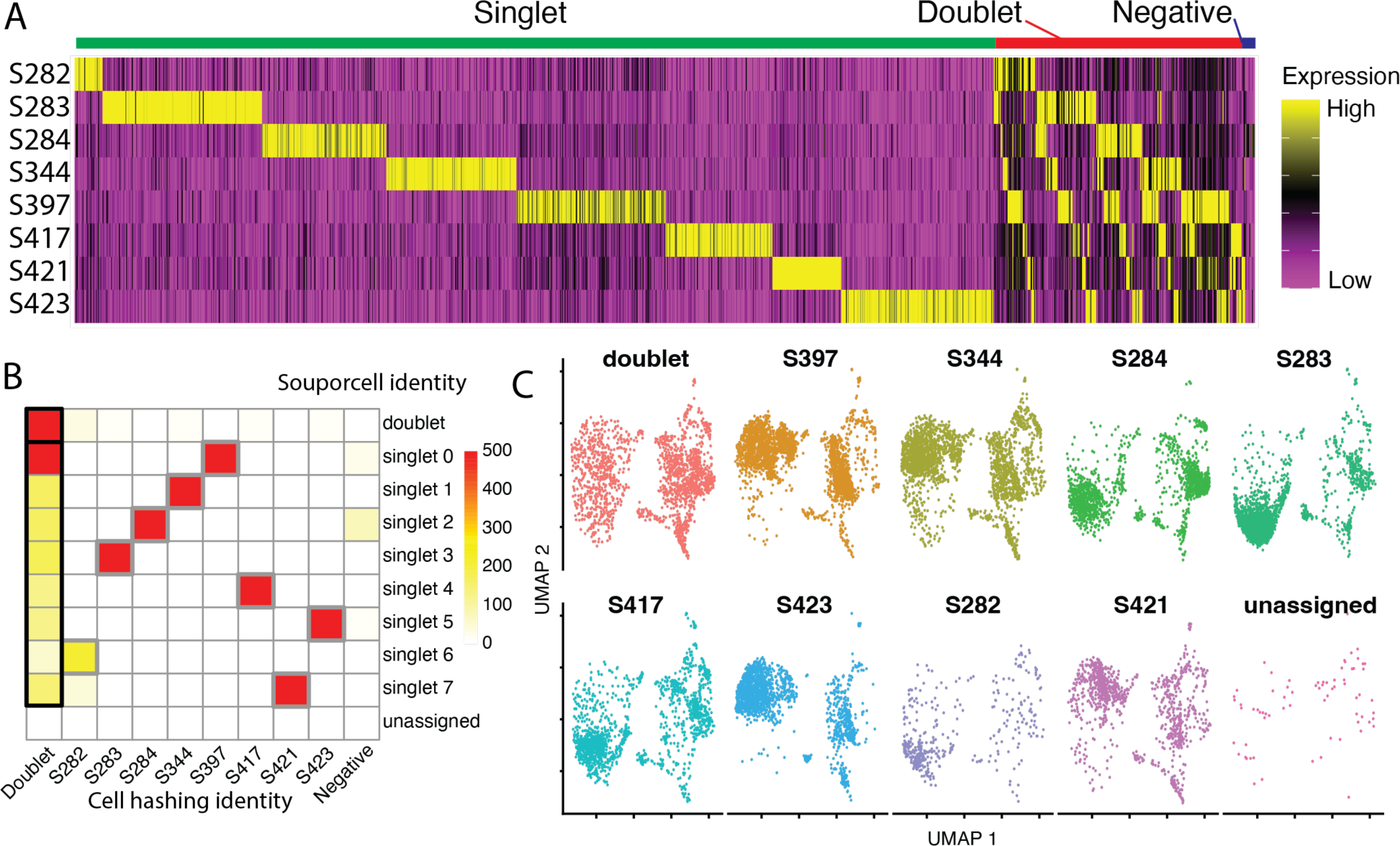
The hybrid strategy combines cell hashing technology and genomic signature method and achieves higher recovery rate and calling accuracy on a real-world immune cell dataset. (**A**) Expression of eight cell hashtags on dataset 8pool-CA. Singlet, doublet, and negative groups are indicated on the top of the heatmap. (**B**) A heatmap of correlation between cell hashing demultiplexing and SNP-based demultiplexing on all cells of dataset 8pool-CA. True doublets and false doublets were highlighted in thick black border, and genotype cluster-cell hashing pairs were highlighted in thick gray border. (**C**) Cells demultiplexed by this hybrid strategy of dataset 8 and dataset 8pool-CA are visualized on an integrated UMAP individually.

**Table 2.**
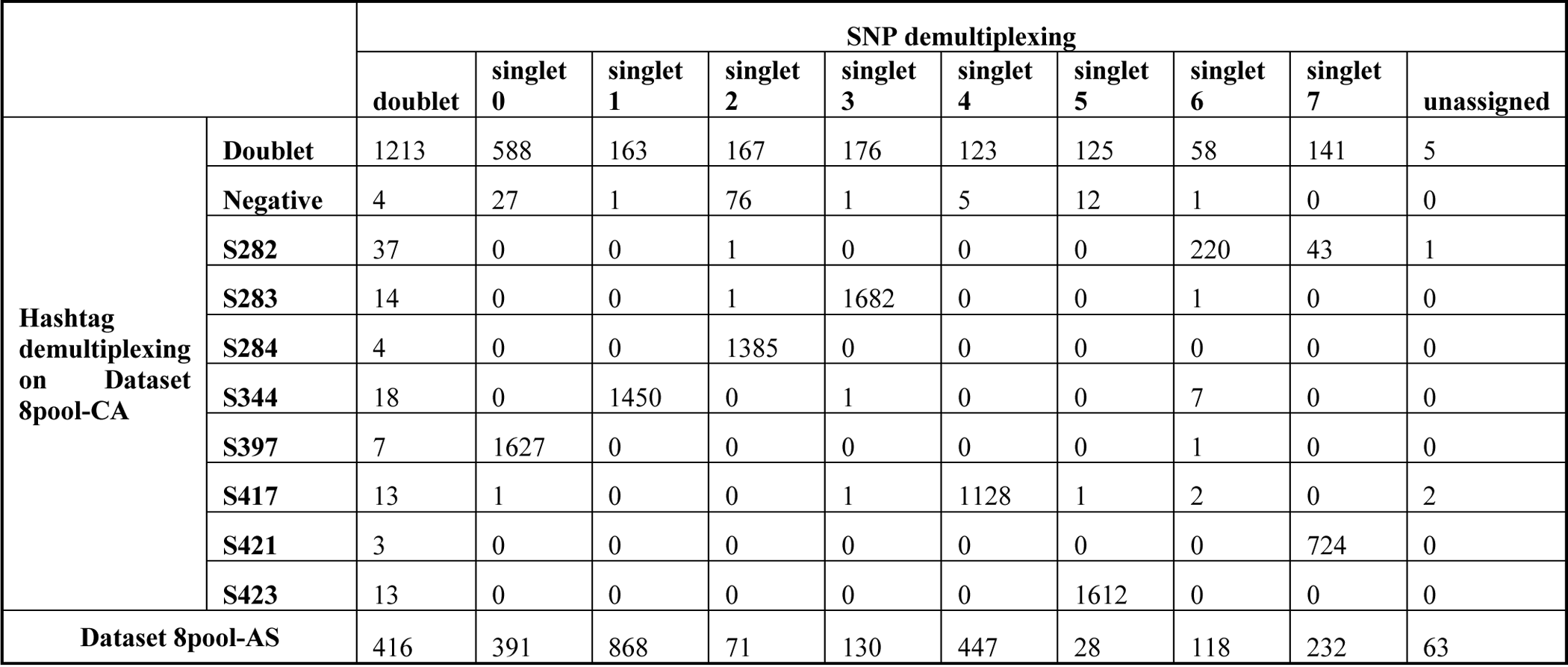
Hybrid demultiplexing on dataset 8pool-CA and 8pool-AS.

Next, we conducted a comprehensive comparison of our hybrid demultiplexing strategy against other cell hashing-based demultiplexing methods, including MULTI_seq, HTOdemux, GMM_Demux, BFF_raw, and BFF_cluster (McGinnis et al., 2019, Hao et al., 2021, Xin et al., 2020, Boggy et al., 2022). Results showed that our hybrid method produced the second highest singlet rate, with the highest generated by BFF_cluster (Table S1). However, it is worth noting that both BFF_raw and BFF_cluster methods failed to identify cells from sample S282, indicating that high recovery rates produced by BFF model may lead to decrease in accuracy when dealing with specific dataset.

In addition to improving cell recovery, this hybrid strategy can also decrease the experimental cost for processing samples with large number of cells. As we mentioned, cells from 8pool-AS and 8pool-CA shared the same donors and SNP profiles, so we ran Souporcell on all cells together to demultiplex cells from 8pool-AS that were not labeled with hashtags. Results showed that we identified 2,285 cells in eight singlet clusters (82.67%), 416 cells in doublet cluster (15.05%), and 63 cells in negative cluster (2.28%) (Table 2), and our hybrid strategy achieved an overall cell recovery rate of 88.42% (13,837 out of 15,650) with all cells from 8pool-AS and 8pool-CA combined (Table 2). Therefore, when processing large scale samples with hybrid strategy, only a small aliquot of cells from each donor needs to be stained with hashtag to link donor identity with SNP profile, and all cells can be efficiently demultiplexed via genetic variation-based clustering. This unique feature not only enhances experimental flexibility, but also reduces consumption of hashtag antibodies per sample without sacrificing performance. Together, compared with single-modal cell hashing method, our hybrid demultiplexing strategy demonstrates enhancement in both cell recovery rate and cost-effectiveness.

### The hybrid demultiplexing strategy is more robust and reliable than single-modal methods

Another important strength of our hybrid strategy is its ability to deliver robust and reliable results, largely independent of hashtag staining quality or performance of the demultiplexing algorithm. For instance, even when GMM_Demux only identified 39.82% of cells in 8pool-CA as singlet, our hybrid strategy still correlated the cell hashing groups generated by GMM_Demux with genotype clusters and accurately determined sample identity of almost 90% of the cells (Table S5). In another extreme case, where two BFF models failed to identify cells from sample S282, our hybrid strategy successfully correlated the incomplete cell hashing groups identified by the BFF models with genotype clusters, and eventually revealed that cluster of singlet6 represented cells from sample S282, as they were not labeled with any other samples’ hashtags (Table S6, S7). Therefore, our hybrid strategy allows up to one unlabeled sample that’s either due to experimental design or failure in cell hashing (N samples with N-1 labeled with hashtags). Unsurprisingly, for the other two methods that generated high quality cell hashing results, our hybrid strategy seamlessly correlated genotype clusters with cell hashing groups (Table S8, S9). Thus, our hybrid strategy produces reliable results as long as there are sufficient single hashtag-labeled cells within each genotype cluster.

Although genetic variation-based methods are known to be highly accurate, we found that Souporcell can generate incorrect genotype clusters in dataset with high doublet rate (Figure 4-A). When we applied our hybrid strategy to dataset 9pool-CA with T and B cells from nine donors, we observed a very poor correlation in singlets between HTOreader and Souporcell (Figure 4-D, E, Table S10). After further analysis, we identified transcriptional clusters consisting of doublets expressing both B cell (CD19, MS4A1) and T cell (CD3E, IL7R) markers (Figure 4-B). Besides these B&T doublets, HTOreader revealed that there were more same-cell-type doublets in other clusters expressing just B cell or T cell markers (Figure 4-C). Therefore, we suspected that the unexpectedly high proportion of true doublets in this dataset might interfere with the unsupervised clustering process of SNP-based methods, making Souporcell unable to accurately generate genotype clusters. To test our hypothesis, we removed cell clusters expressing both B and T cell markers and found that the correlation increased to 54.52% (Figure 4-F, Table S11). After we removed all doublets assigned by HTOreader (Figure 4-D), correlation reached 96.06% and demultiplexing results went back to normal (Figure 4-G, Table S12). These results highlight the risk associated with genetic variant-based methods that use unsupervised clustering algorithms, especially when applied to dataset enriched with true doublets. By correlating results from nucleotide barcode and natural genetic variation-based methods, our hybrid strategy allows two methods to complement each other and provide self-correction against potential errors, making it more robust and reliable than single-modal methods.

**Figure 4.**
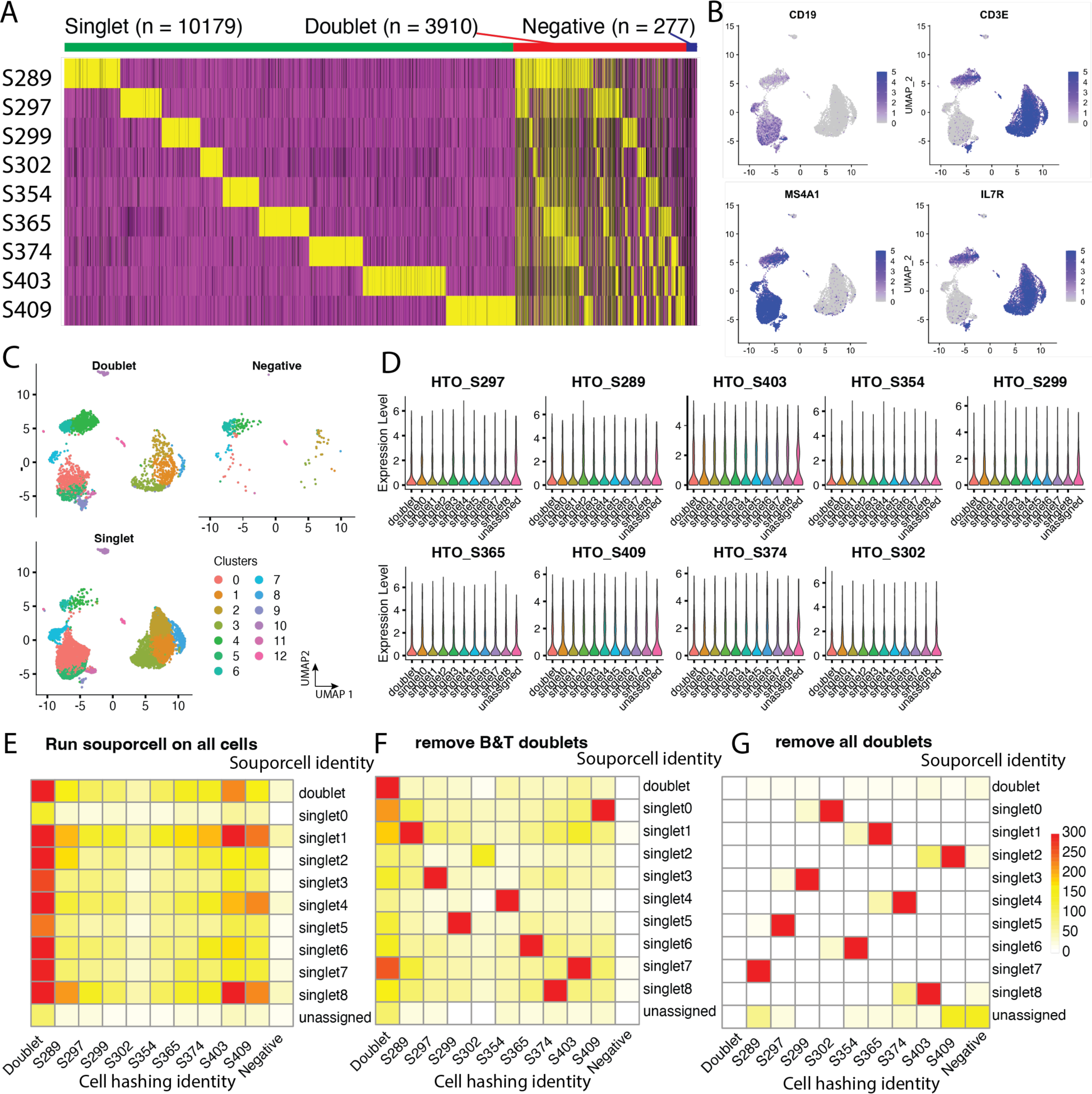
Hybrid demultiplexing strategy corrects potential error of SNP-based method. (**A**) Expression of nine cell hashtags on dataset 9pool-CA. Singlet, doublet, and negative groups are indicated on the top of the heatmap. (**B**) Expression of two B cell gene markers (CD19 and MS4A1) and two T cell gene markers (CD3E and IL7R) visualized on a UMAP embedding of carrier cells of dataset 9pool-CA. (**C**) Doublets, negative cells, and singlets identified by cell hashing method are visualized on a UMAP individually. Cells are colored by transcriptome clusters. (**D**) Expression of nine hashtags on genotype clusters generated by SNP-based method. (**E**) A heatmap of correlation between cell hashing demultiplexing and SNP-based demultiplexing on all cells of dataset 9pool-CA. (**F**) A heatmap of correlation between cell hashing demultiplexing and SNP-based demultiplexing on all cells of dataset 9pool-CA after remove B&T doublets (cells that express both B and T cell markers). (**G**) A heatmap of correlation between cell hashing demultiplexing and SNP-based demultiplexing on all cells of dataset 9pool-CA after all potential doublets (cells that express more than one hashtag).

## Discussion

Over the last few years, sample super loading and demultiplexing strategies have been extensively studied to improve the quality, magnitude, and economy of single-cell experiments. To date, there are two major demultiplexing solutions, nucleotide barcode-based methods that link sequences of unique oligonucleotide tags to sample identity, and natural genetic variation-based methods that identify donor identity by natural single nucleotide polymorphisms (SNPs). Among them, cell hashing that derives from CITE-seq technology has been one of the most commonly used demultiplexing approaches due to its compatibility and simplicity. However, cell hashing has several disadvantages that remain to be improved, including difficulties in distinguishing true positive from background signal, unpredictable false negative and false doublet rates, and high reagent cost when staining large number of cells. Here, we propose a hybrid demultiplexing strategy that integrates results of cell hashing and SNP clustering. By testing it on multiple real-word datasets, we demonstrate it consistently improves accuracy, cell recovery rate, cost-effectiveness and robustness of cell hashing.

R and Python are two major coding languages used in single-cell analysis. In R community, Seurat package is the most widely used pipeline for downstream analysis due to its versatility, rich documentations, usability, and compatibility with multi-omics. As we demonstrated above, the default demultiplexing algorithm for cell hashing in Seurat performs poorly in terms of cell recovery for datasets with imbalanced sample sizes (Fig 1B-D). We found that Seurat often generates inaccurate cutoffs for each hashtag when staining quality or number of cells varies across samples, causing a large number of cells with low hashtag signal incorrectly classified as negative. To solve this, we developed HTOreader for R users, a toolkit that improves cut-off calling accuracy when demultiplexing cell hashing data with Seurat. Compared to Seurat, we demonstrate that HTOreader delivers much higher accuracy and cell recovery rate on real-world datasets with imbalanced sample sizes.

One major caveat of cell hashing is that the cell recovery rate decreases as the number of hashtags being used increases due to the accumulation of false doublets. Based on our experience, it is completely normal to lose 25-40% of cells when using more than 8 hashtags at once. In addition, poor staining quality of certain hashtag that occurs unpredictably can further decrease cell recovery by enrichment of false negatives. Recently, advances in bioinformatics allow users to cluster cells from each human donor according to their SNP profiles. This genotype clustering-based demultiplexing approach is highly accurate and can recovery up to 90% of cells. However, it is not widely used because it can only be used with human donors and requires users to generate SNP references for each donor via genomic DNA sequencing or bulk RNA-seq to link donor identities to genotype clusters. Therefore, we developed a hybrid demultiplexing strategy that integrates results of cell hashing and genotype clustering so that the two approaches can validate and complement each other. Our results showed that this hybrid strategy consistently increases cell recovery to 90% regardless of hashtag numbers or staining quality. Since only a small fraction of singlet determined by cell hashing is sufficient to link donor identities with each genotype cluster, the reagent cost of hashtag antibodies can be greatly decreased, and generation of SNP references is no longer needed. Additionally, this approach is extremely useful to simultaneously demultiplex multiple pooled samples from the same group of donors (such as longitudinal studies) because only cells from one sample need to be stained with hashtags for obtaining donor identities and cells from all samples can be demultiplexed by genotype clustering (e.g., 9pool-CA+9pool-AS, 8pool-CA+8pool-AS). We also found that this strategy corrects errors caused by one of the two methods and allows users to use N hashtags to demultiplex samples with N+1 donors. To conclude, our hybrid strategy provides a neat solution that improves performance, robustness, and cost-effectiveness of the existing single-modal methods, such as cell hashing and genotype clustering.

Sample super-loading and demultiplexing will remain important topics for single cell studies. Recently, the 10x Genomics 3’ CellPlex, a lipid-based cell hashing technology, provides a promising alternative that might solve some of the issues of antibody-based cell hashing approach we mentioned in this article. So far, CellPlex is only available for 3’ sequencing, which is not compatible with single cell immune profiling and some other applications. However, as single cell sequencing becomes more common, more demultiplexing techniques are going to be available in the near future to greatly benefit biological and medical research.

## Material and Methods

### Datasets

#### Dataset 1

A single-cell dataset from human peripheral blood mononuclear cells (referred to as Stoeckius-2018 hereafter). This dataset is comprised of 8 individual donors that are uniquely labeled by 8 cell hashtags. This dataset has been published with cell hashing original paper(Stoeckius et al., 2018). Dataset is available from https://www.dropbox.com/sh/ntc33ium7cg1za1/AAD_8XIDmu4F7lJ-5sp-rGFYa?dl=0. More details of this dataset can also be found from Seurat website: https://satijalab.org/seurat/articles/hashing_vignette.html.

#### Dataset 2

A novel single-cell dataset generated for this paper, labeled by subject ID, 3V007 (referred to as 3V007 hereafter). In this dataset, mRNA, B Cell Receptor (BCR) repertoire, surface protein expression (CD27 and CD79b), and binding affinity of 14 antigen-probes, including HA proteins of several endemic influenza strains, were measured. Two groups of cells (from same donor), antigen-specific B cell (influenza HA specific) and carrier cells (T cells and B cells) were uniquely labeled by 2 cell hashtags, hashtag1 and hashtag2, respectively. These three hashtags were also sequenced with those antigen-probes. This dataset has been uploaded to GEO database (see data availability statement for details).

#### Dataset 3

A novel single-cell dataset generated for this paper, labeled by subject ID, S414 (referred to as S414 hereafter). In this dataset, mRNA, B Cell Receptor (BCR) repertoire, surface protein expression (CD27 and CD79b), and binding affinity of 8 antigen-probes, including HA proteins of several endemic influenza strains, were measured. Four groups of cells (from same donor) were uniquely labeled by 4 cell hashtags, hashtag3, hashtag4, hashtag5, and hashtag6. We splitted carrier cells (T cells with a few B cells) into two groups, labeled them with hashtag3 and hashtag4; and splitted antigen-specific B cells into two groups, labeled them with hashtag5 and hashtag6. These four hashtags were also sequenced with those antigen-probes. This dataset has been uploaded to GEO database (see data availability statement for details).

#### Dataset 4

This dataset (dataset name is R125, referred to as R125 hereafter) is from a published single-cell dataset from a previous publication(Dugan et al., 2021). In this dataset, mRNA, B Cell Receptor (BCR) repertoire, and binding affinity of 17 antigen-probes, including Spike, NP, ORF8 and RBD protein of endemic and pandemic COVID strain, HA protein of influenza virus, and interferon alpha and omega, were measured. Cells from 3 individual human donors were uniquely labeled by 3 cell hashtags, hashtag1-R1, hashtag2-R2, and hashtag3-R5. These three hashtags were also sequenced with these antigen-probes. This dataset (R125) is available from Mendeley Data: https://doi.org/10.17632/3jdywv5jrv.3.

#### Dataset 5

This dataset (dataset name is R6, referred to as R6 hereafter) is from a published single-cell dataset from a previous publication(Dugan et al., 2021). In this dataset, mRNA, B Cell Receptor (BCR) repertoire, and binding affinity of 17 antigen-probes, including Spike, NP, ORF8 and RBD protein of endemic and pandemic COVID strain, HA protein of influenza virus, and interferon alpha and omega, were measured. Cells from 2 time points of an individual human donor were uniquely labeled by 2 cell hashtags, hashtag3-early and hashtag4-late. These three hashtags were also sequenced with these antigen-probes. This dataset (R6) is available from Mendeley Data: https://doi.org/10.17632/3jdywv5jrv.3.

#### Dataset 6

A novel single-cell dataset generated for this paper, labeled as 9pool-CA (CA is short for carrier, referred to as 9pool-CA hereafter). In this dataset, we sorted B cells and T cells from nine subjects and pooled them together. mRNA, and surface protein expression panel were measured. Cells from nine individual human donors were uniquely labeled by nine cell hashtags, hashtag1 to hashtag9. These nine hashtags were also sequenced with those surface proteins. This dataset has been uploaded to GEO database (see data availability statement for details).

#### Dataset 7

A novel single-cell dataset generated for this paper, labeled as 9pool-AS (AS is short for antigen-specific, referred to as 9pool-AS hereafter). In this dataset, we sorted antigen-specific B cells from nine human donors (as same as dataset 6) and pooled them together. mRNA, B Cell Receptor (BCR) repertoire, surface protein expression (CD27 and CD79b), and binding affinity of 18 antigen-probes, including HA proteins of several endemic influenza strains, were measured. This dataset has been uploaded to GEO database (see data availability statement for details).

#### Dataset 8

A novel single-cell dataset generated for this paper, labeled as 8pool-AS (referred to as 8pool-AS hereafter). In this dataset, we sorted antigen-specific B cells from eight subjects and pooled them together. mRNA, B Cell Receptor (BCR) repertoire, surface protein expression (CD27 and CD79b), and binding affinity of several antigen-probes, including HA proteins of several endemic influenza strains, were measured. This dataset has been uploaded to GEO database (see data availability statement for details).

#### Dataset 9

A novel single-cell dataset generated for this paper, labeled as 8pool-CA (referred to as 8pool-CA hereafter). In this dataset, we sorted B cells and T cells from eight subjects (as same as dataset 8) and pooled them together. mRNA, and surface protein expression panel were measured. Cells from eight individual human donors were uniquely labeled by eight cell hashtags, hashtag1 to hashtag8. These eight hashtags were also sequenced with those surface proteins. This dataset has been uploaded to GEO database (see data availability statement for details).

### Cell hashing demultiplexing methods

We developed and introduced an improved demultiplexing approach for single-cell cell hashing, called HTOreader. To accurately determine the hashtag identity for each individual cell, we developed a cutoff calling method that precisely distinguishes true positive from background. Specifically, the distributions of normalized counts for each hashtag were first fitted into two Gaussian distributions, representing background and true positive groups. Then a cutoff value that distinguishes the two groups was calculated based on means and standard divisions of these two Gaussian distributions. Finally, the identity of each individual cell was determined according to the hashtags they were positive of.

#### Data normalization

Two normalization methods: Centered Log-Ratio (CLR) and Log (*log*1*p*) normalization are available. CLR method is more common in normalization of CITE-seq protein expression and hashtags(Aitchison, 1982, Hao et al., 2021). For a given raw counts vector *W* of a hashtag, the CLR normalization will be:

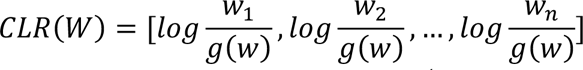

Where *n* is the length of vector *W*, and 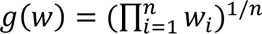 denotes the geometric mean of *W*. The conventional Log normalization works well in some datasets. We use *log*1*p* to avoid the undefined log[0]. For a given raw counts vector *W* of a hashtag, the Log normalization will be:

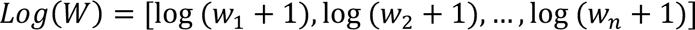

#### Mixture modeling

Mixture modeling has been extensively used in single-cell data pre-processing, such as estimation of the drop-out rate, determination of effective sequencing depth and amplification noise(Fan et al., 2016, Kharchenko et al., 2014). We adopted an mixture modeling approach implemented in the Flexmix package to fit two Gaussian distributions from a vector of normalized hashtag counts(Leisch, 2004). In this step, we fit normalized data of each hashtag into two Gaussian distributions indicating one positive group, representing background and true positive groups, and calculate the means and standard deviations of these two groups respectively.

#### Cutoff determination

For two Gaussian distributions 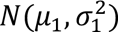 and 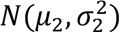, 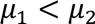, we determine the cutoff to distinguish true positive and background using the following equation:

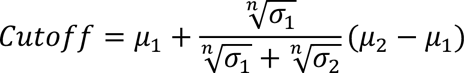

Where *n* is the rank of the model, the recommended rank is 2 in most cases. Please see the supplementary material for details.

#### Sample identity assignment

For each cell, we assign their sample identities based on their binding status of each hashtag(Hao et al., 2021). If a cell is deemed positive for only one hashtag, it will be labeled as a singlet for that corresponding hashtag; if it’s deemed positive to multiple hashtags, it will be labeled as doublet; if it’s deemed background for all hashtags, it will be labeled as negative. Sample identities of every cell labeled as singlet will be assigned according to their hashtag identities.

### Genomic signature demultiplexing

Genomic signature demultiplexing method is an essential part of this hybrid demultiplexing strategy, and to date many computational methods are available, for example demuxlet and souporcell. In this paper, we applied souporcell, which has been widely used in the community, onto our strategy to demonstrate the effectiveness of this workflow. As most SNP-based demultiplexing methods do, souporcell aligns all short reads against a reference genome to get the SNPs for each cell, then groups cells into multiple clusters according to their genotypes (SNP signatures) using an unsupervised learning algorithm. The number of genotype clusters is pre-defined by users according to the number of subjects in the pooled sample. For a sample pooled from N subjects (individual human donors), there will be N+2 distinct genotype clusters identified, in which one “doublet” cluster containing cells fit in more than one genotypes, one “negative” cluster containing cells whose SNP signatures are not sufficient therefore cannot be fit into any genotype, and N singlet clusters indicating cells from N individual donors.

### Cell hashing demultiplexing comparision

We conducted demultiplexing using all existing methods, including MULTI-seq, GMM-Demux, HTOdemux, BFF_raw, and BFF_cluster, utilizing the R package cellhashR (version 1.0.3) in R 4.2.2. Additionally, we ran HTOreader using the R package HTOreader (version 0.1.0) in R 4.2.2. For cellhashR, all parameters were set to default, while for HTOreader, we used the ‘log’ option for the ‘method’ parameter with the remaining parameters set to their default values.

### Flow cytometry staining and cell sorting

Flow staining and cell sorting were performed as previously described (Dugan et al., 2021). Briefly, human PBMCs were thawed in 10% FBS RPMI1640 medium and enriched by negative selection using a pan-B cell isolation kit according the manufacturer’s instruction (StemCell, Cat#. 19554) prior to staining with the following antibodies and flurorescently oligonucleotide-labeled streptavidin-antigen tetramers (Biolegend) : anti-huCD19-PE-Cy7, anti-huCD3-BB515, anti-huCD4-BB515, anti-huIgD-BB515, TotalSeq-C anti-human hashtag antibodies, antigen-PE or- APC, and at 4 degree for 30 mins. Cells were subsequently washed three times with 2% FBS PBS buffer supplemented with 2mM D-biotin. Finally, cells were adjusted at a maximum of 2 million cells per ml in washing buffer, stained with DAPI and subjected to sorting by either MACSQuantTyto (Miltenyi) or BD Melody (BD). Cells that were viable/CD19^+^/antigen-PE^+^ and antigen-APC^+^ or viable/CD4^+^ were sorted for downstream 10X Genomics processing.

### 10X Genomics libraries construction and Next Generation Sequencing

5’ gene expression, VDJ and surface protein feature libraries were prepared using the 10X genomics platform as per the manufacturer’s instructions (Chromium Next GEM Single Cell 5’ (HT) Reagent Kits v2 (Dual Index)). Three libraries were quantified by real-time quantitative PCR using KAPA Library Quanitification Kits (Roche) and pooled at recommended ratio and sequenced using NextSeq1000 (Illumina) with 26 cycles for read 1, 10 cycles for i7/i5 index, 150 cycles for read 2.

## Data Availability Statement

All single-cell datasets generated for this study have been uploaded into GEO database. The GEO accession number is GSE230810.

## Code Availability Statement

We have deposited the source code of the R package we developed in this study to GitHub: https://github.com/WilsonImmunologyLab/HTOreader.

## Acknowledgements

This project was funded in part by the National Institute of Allergy and Infectious Diseases (NIAID; National Institutes of Health grant numbers U19AI082724 (PCW), U19AI109946 (PCW), U19AI057266 (PCW), the NIAID Centers of Excellence for Influenza Research and Surveillance (CEIRS) grant number HHSN272201400005C (PCW), and the NIAID Centers of Excellence for Influenza Research and Response (CEIRR) grant number 75N93019R00028 (PCW). This work was also partially supported by the NIAID Collaborative Influenza Vaccine Innovation Centers (CIVIC; 75N93019C00051). In addition, this work was also partially funded by the NIAID Centers of Excellence for Influenza Research and Surveillance (CEIRS, contract # HHSN272201400008C), the NIAID Centers of Excellence for Influenza Research and Response (CEIRR, contract# 75N93021C00014) and by anonymous donors. JJCM was supported by a CIHR Banting Postdoctoral Fellowship (BPF-186528).

## Author Contribution

LL designed the model and workflow, implemented the package, performed computational analyses of single-cell data, and wrote the manuscript. JS and YF collected samples, designed the workflow, performed experiments, and wrote the manuscript.SC and JJCM collected samples, helped in workflow design, and revised the manuscript. PCW supervised the work, designed the workflow, and wrote the manuscript.

## Declaration of interests

The authors declare no competing interests.

## Notes

### Competing Interest Statement

The authors have declared no competing interest.

### Summary of Updates

We re-organized and revised the introduction, results, and discussion sections according to reviewers' comments. We added comparisons with existing methods and made the manuscript more concise and focused.

