## Supplementary material for "A hybrid demultiplexing strategy that improves performance and robustness of cell hashing": Fig S1, and S2, Table S1 - S12

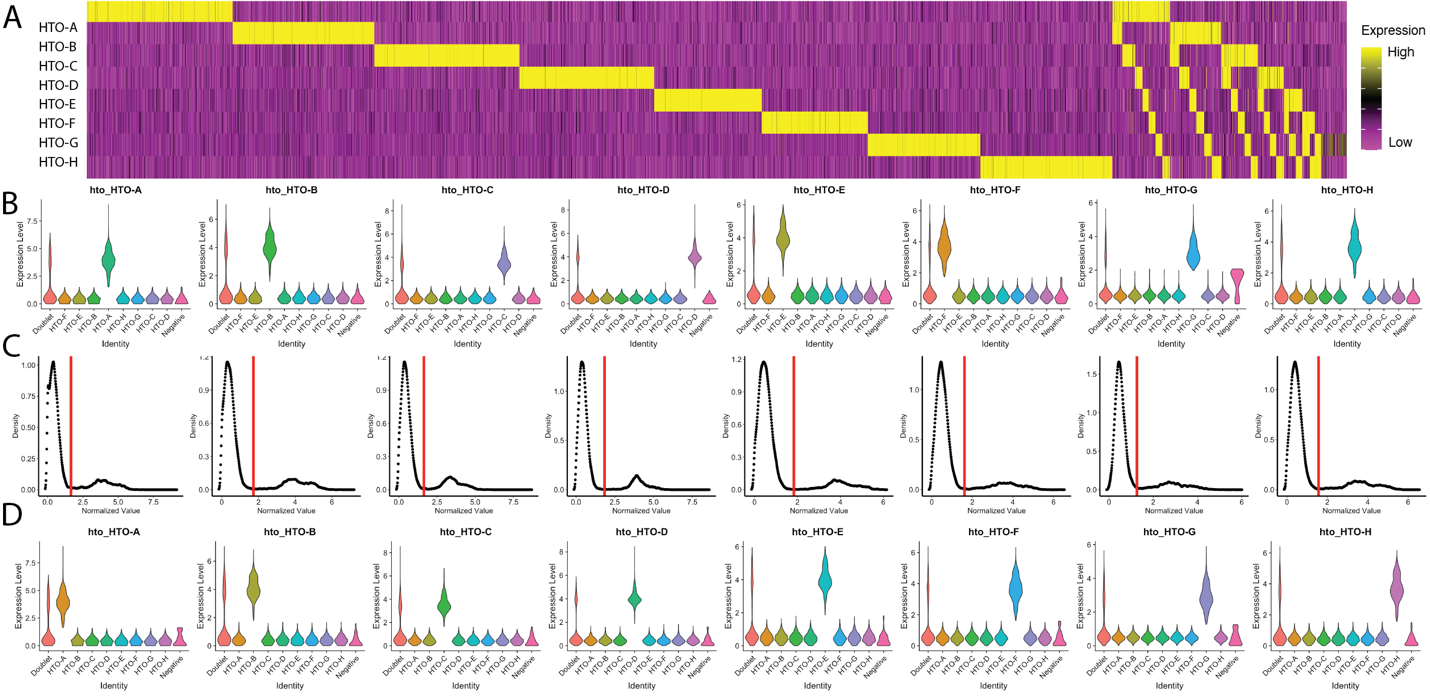

**Figure S1**. Comparing cell demultiplexing on a public benchmark dataset using both Seurat and HTOreader. (**A**) expression heatmap of all eight hashtags ordered by Seurat labels. (**B**) violin plot of expression of all eight hashtags on Seurat groups. (**C**) cutoffs determined by HTOreader based on density of normalized values for all eight hashtags. (**D**) violin plot of expression of all eight hashtags on HTOreader groups.

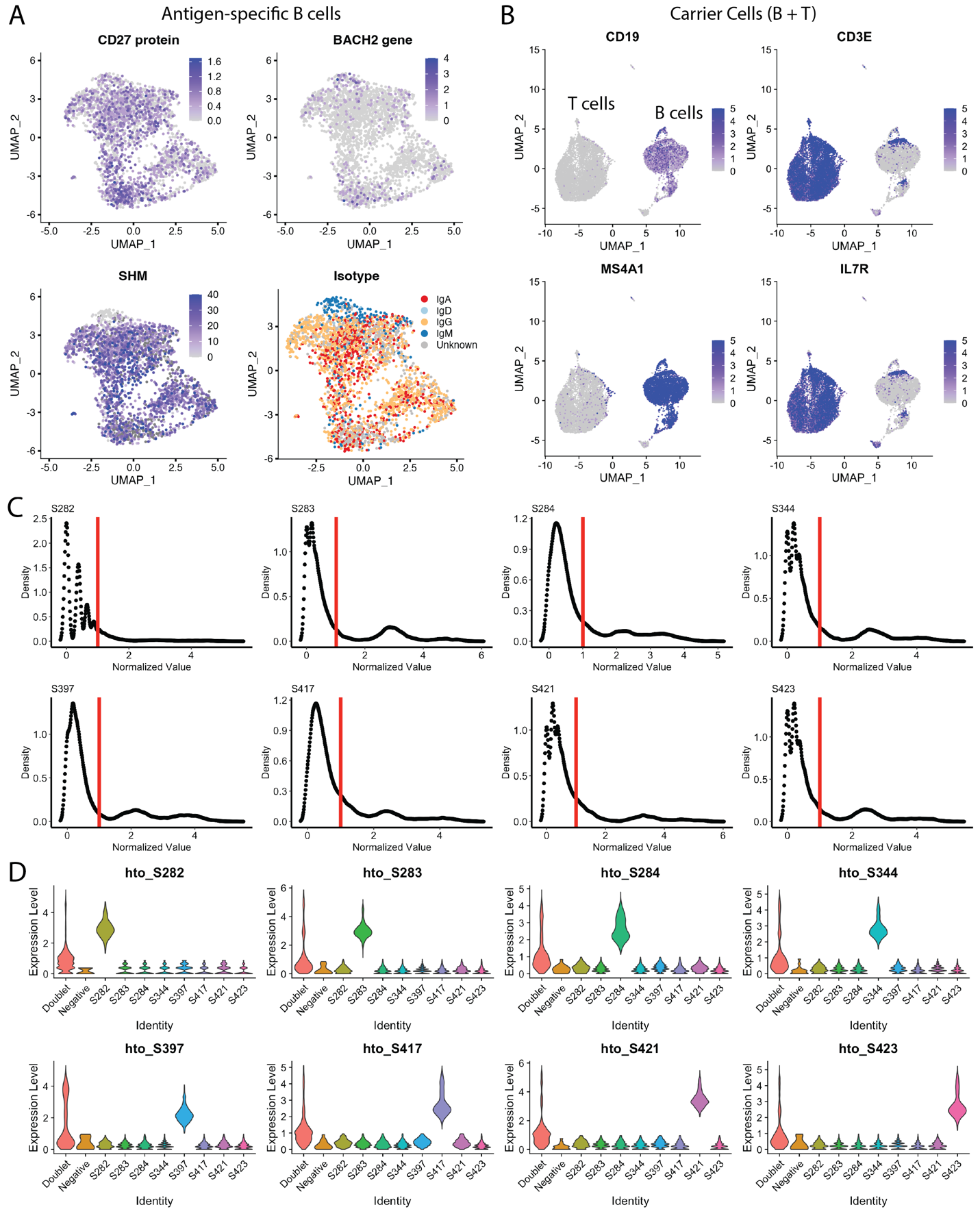

**Figure S2**. Cell demultiplexing on a B cells and T cells pooled dataset (8pool-CA and 8pool-AS) using hybrid method. (**A**) expression of CD27 protein, expression of BACH2 gene, Somatic hypermutation (SHM) of heavy chain, and isotype of BCR repertoire visualized on a UMAP embedding of antigen-specific B cells of 8pool-AS. (**B**) Expression of two B cell gene markers (CD19 and MS4A1) and two T cell gene markers (CD3E and IL7R) visualized on a UMAP embedding of carrier cells of 8pool-CA. T cell and B cell clusters are roughly indicated by labels (**C**) Cutoffs determined by HTOreader based on density of normalized values for all eight hashtags on 8pool-CA. (**D**) Violin plot of expression of all eight hashtags on HTOreader demultiplexing groups on 8pool-CA.

**Table S1**. Cell hashing demultiplexing of dataset 8pool-CA using existing methods. Number of total singlets and the singlet rate were highlighted in bold.

|  | **Multi-seq** | **HTOdemux** | **GMM_Demux** | **BFF_raw** | **BFF_cluster** | **HTOreader** |
| --- | --- | --- | --- | --- | --- | --- |
| Doublet | 2120 | 3584 | 7747 | 1704 | 653 | 2759 |
| Negative | 159 | 20 | 16 | 538 | 518 | 127 |
| S282 | 308 | 278 | 321 | 0 | 0 | 302 |
| S283 | 1740 | 1567 | 755 | 1794 | 1961 | 1698 |
| S284 | 1545 | 1372 | 749 | 1525 | 1670 | 1389 |
| S344 | 1548 | 1354 | 751 | 1583 | 1729 | 1476 |
| S397 | 1855 | 1378 | 541 | 2080 | 2313 | 1635 |
| S417 | 1197 | 1093 | 682 | 1196 | 1331 | 1148 |
| S421 | 779 | 661 | 293 | 791 | 879 | 727 |
| S423 | 1635 | 1579 | 1031 | 1675 | 1832 | 1625 |
| **Singlet** | **10607** | **9282** | **5123** | **10644** | **11715** | **10000** |
| **Singlet rate** | **82.44%** | **72.14%** | **39.82%** | **82.73%** | **91.05%** | **77.72%** |

**Table S2**. Cell hashing demultiplexing of dataset S414 using existing methods.

|  | **Multi-seq** | **HTOdemux** | **GMM_demux** | **BFF_raw** | **BFF_cluster** | **HTOreader** |
| --- | --- | --- | --- | --- | --- | --- |
| Doublet | 696 | 1052 | 928 | 772 | 781 | 967 |
| hashtag3 | 2614 | 2305 | 2548 | 2612 | 2595 | 2527 |
| hashtag4 | 2173 | 2067 | 2123 | 2143 | 2129 | 2115 |
| hashtag5 | 802 | 840 | 802 | 817 | 810 | 798 |
| hashtag6 | 738 | 775 | 725 | 747 | 739 | 723 |
| Negative | 250 | 234 | 147 | 182 | 219 | 143 |
| Singlet | 6327 | 5987 | 6198 | 6319 | 6273 | 6163 |
| Singlet rate | 86.99% | 82.32% | 85.22% | 86.88% | 86.25% | 84.74% |

**Table S3**. Cell hashing demultiplexing of dataset 3V007 using existing methods.

|  | **Multi-seq** | **HTOdemux** | **GMM_demux** | **BFF_raw** | **BFF_cluster** | **HTOreader** |
| --- | --- | --- | --- | --- | --- | --- |
| Doublet | 1032 | 984 | 1152 | 1099 | 1149 | 1135 |
| hashtag1 | 4304 | 3750 | 4368 | 4372 | 4320 | 4355 |
| hashtag2 | 4598 | 4746 | 4478 | 4521 | 4469 | 4505 |
| Negative | 65 | 519 | 1 | 7 | 61 | 4 |
| Singlet | 8902 | 8496 | 8846 | 8893 | 8789 | 8860 |
| Singlet rate | 89.03% | 84.97% | 88.47% | 88.94% | 87.90% | 88.61% |

**Table S4**. Cell hashing demultiplexing of dataset R125 using existing methods.

|  | **Multi-seq** | **HTOdemux** | **GMM_demux** | **BFF_raw** | **BFF_cluster** | **HTOreader** |
| --- | --- | --- | --- | --- | --- | --- |
| Doublet | 39 | 130 | 66 | 52 | 42 | 78 |
| Hashtag1-R1 | 173 | 168 | 155 | 150 | 171 | 160 |
| Hashtag2-R2 | 38 | 268 | 444 | 416 | 528 | 437 |
| Hashtag3-R5 | 239 | 224 | 218 | 216 | 241 | 215 |
| Negative | 500 | 199 | 106 | 155 | 7 | 99 |
| Singlet | 450 | 660 | 817 | 782 | 940 | 812 |
| Singlet rate | 45.50% | 66.73% | 82.61% | 79.07% | 95.05% | 82.10% |

**Table S5**. Correlation between Souporcell genotype clusters and GMM_Demux method of dataset 8pool-CA.

|  | doublet | singlet 0 | singlet 1 | singlet 2 | singlet 3 | singlet 4 | singlet 5 | singlet 6 | singlet 7 | unassigned |
| --- | --- | --- | --- | --- | --- | --- | --- | --- | --- | --- |
| Doublet | 1250 | 1696 | 871 | 880 | 1108 | 580 | 719 | 66 | 571 | 6 |
| Negative | 3 | 3 | 1 | 2 | 2 | 2 | 3 | 0 | 0 | 0 |
| S282 | 45 | 6 | 0 | 2 | 0 | 0 | 3 | 220 | 44 | 1 |
| S283 | 5 | 0 | 0 | 0 | 750 | 0 | 0 | 0 | 0 | 0 |
| S284 | 2 | 0 | 0 | 746 | 0 | 0 | 0 | 1 | 0 | 0 |
| S344 | 5 | 0 | 742 | 0 | 1 | 0 | 0 | 3 | 0 | 0 |
| S397 | 3 | 538 | 0 | 0 | 0 | 0 | 0 | 0 | 0 | 0 |
| S417 | 7 | 0 | 0 | 0 | 0 | 674 | 0 | 0 | 0 | 1 |
| S421 | 0 | 0 | 0 | 0 | 0 | 0 | 0 | 0 | 293 | 0 |
| S423 | 6 | 0 | 0 | 0 | 0 | 0 | 1025 | 0 | 0 | 0 |

**Table S6**. Correlation between Souporcell genotype clusters and BFF_raw method of dataset 8pool-CA.

|  | doublet | singlet 0 | singlet 1 | singlet 2 | singlet 3 | singlet 4 | singlet 5 | singlet 6 | singlet 7 | unassigned |
| --- | --- | --- | --- | --- | --- | --- | --- | --- | --- | --- |
| Doublet | 1090 | 145 | 68 | 80 | 93 | 61 | 73 | 26 | 63 | 5 |
| Negative | 59 | 45 | 8 | 55 | 11 | 37 | 37 | 227 | 58 | 1 |
| S282 | 0 | 0 | 0 | 0 | 0 | 0 | 0 | 0 | 0 | 0 |
| S283 | 33 | 0 | 1 | 1 | 1755 | 0 | 0 | 4 | 0 | 0 |
| S284 | 28 | 0 | 0 | 1494 | 0 | 1 | 0 | 1 | 1 | 0 |
| S344 | 33 | 0 | 1537 | 0 | 1 | 2 | 0 | 10 | 0 | 0 |
| S397 | 18 | 2053 | 0 | 0 | 0 | 0 | 0 | 7 | 2 | 0 |
| S417 | 30 | 0 | 0 | 0 | 1 | 1155 | 1 | 7 | 0 | 2 |
| S421 | 5 | 0 | 0 | 0 | 0 | 0 | 0 | 2 | 784 | 0 |
| S423 | 30 | 0 | 0 | 0 | 0 | 0 | 1639 | 6 | 0 | 0 |

**Table S7**. Correlation between Souporcell genotype clusters and BFF_cluster method of dataset 8pool-CA.

|  | doublet | singlet 0 | singlet 1 | singlet 2 | singlet 3 | singlet 4 | singlet 5 | singlet 6 | singlet 7 | unassigned |
| --- | --- | --- | --- | --- | --- | --- | --- | --- | --- | --- |
| Doublet | 475 | 28 | 15 | 26 | 34 | 16 | 26 | 14 | 17 | 2 |
| Negative | 59 | 45 | 6 | 55 | 11 | 31 | 29 | 227 | 54 | 1 |
| S282 | 0 | 0 | 0 | 0 | 0 | 0 | 0 | 0 | 0 | 0 |
| S283 | 139 | 2 | 5 | 2 | 1804 | 3 | 0 | 4 | 1 | 1 |
| S284 | 118 | 5 | 2 | 1536 | 2 | 1 | 0 | 1 | 3 | 2 |
| S344 | 121 | 2 | 1579 | 1 | 4 | 4 | 3 | 15 | 0 | 0 |
| S397 | 135 | 2152 | 3 | 3 | 1 | 2 | 3 | 10 | 4 | 0 |
| S417 | 114 | 2 | 1 | 1 | 3 | 1194 | 2 | 9 | 3 | 2 |
| S421 | 41 | 0 | 1 | 4 | 1 | 2 | 4 | 2 | 824 | 0 |
| S423 | 124 | 7 | 2 | 2 | 1 | 3 | 1683 | 8 | 2 | 0 |

**Table S8**. Correlation between Souporcell genotype clusters and HTOdemux method of dataset 8pool-CA.

|  | doublet | singlet 0 | singlet 1 | singlet 2 | singlet 3 | singlet 4 | singlet 5 | singlet 6 | singlet 7 | unassigned |
| --- | --- | --- | --- | --- | --- | --- | --- | --- | --- | --- |
| Doublet | 1238 | 865 | 281 | 260 | 302 | 176 | 175 | 64 | 218 | 5 |
| Negative | 3 | 5 | 1 | 3 | 1 | 1 | 6 | 0 | 0 | 0 |
| S282 | 30 | 0 | 0 | 0 | 0 | 0 | 0 | 217 | 30 | 1 |
| S283 | 9 | 0 | 0 | 0 | 1557 | 0 | 0 | 1 | 0 | 0 |
| S284 | 4 | 0 | 0 | 1367 | 0 | 0 | 0 | 1 | 0 | 0 |
| S344 | 16 | 0 | 1332 | 0 | 0 | 0 | 0 | 6 | 0 | 0 |
| S397 | 5 | 1373 | 0 | 0 | 0 | 0 | 0 | 0 | 0 | 0 |
| S417 | 9 | 0 | 0 | 0 | 1 | 1079 | 1 | 1 | 0 | 2 |
| S421 | 1 | 0 | 0 | 0 | 0 | 0 | 0 | 0 | 660 | 0 |
| S423 | 11 | 0 | 0 | 0 | 0 | 0 | 1568 | 0 | 0 | 0 |

**Table S9**. Correlation between Souporcell genotype clusters and MULTI_seq method of dataset 8pool-CA.

|  | doublet | singlet 0 | singlet 1 | singlet 2 | singlet 3 | singlet 4 | singlet 5 | singlet 6 | singlet 7 | unassigned |
| --- | --- | --- | --- | --- | --- | --- | --- | --- | --- | --- |
| Doublet | 1182 | 335 | 92 | 92 | 120 | 76 | 82 | 51 | 85 | 5 |
| Negative | 4 | 61 | 5 | 3 | 21 | 5 | 43 | 16 | 1 | 0 |
| S282 | 48 | 0 | 0 | 0 | 0 | 0 | 1 | 211 | 47 | 1 |
| S283 | 20 | 0 | 1 | 0 | 1718 | 0 | 0 | 1 | 0 | 0 |
| S284 | 9 | 0 | 0 | 1535 | 0 | 0 | 0 | 1 | 0 | 0 |
| S344 | 22 | 0 | 1516 | 0 | 1 | 0 | 1 | 8 | 0 | 0 |
| S397 | 8 | 1846 | 0 | 0 | 0 | 0 | 1 | 0 | 0 | 0 |
| S417 | 15 | 1 | 0 | 0 | 1 | 1175 | 1 | 2 | 0 | 2 |
| S421 | 4 | 0 | 0 | 0 | 0 | 0 | 0 | 0 | 775 | 0 |
| S423 | 14 | 0 | 0 | 0 | 0 | 0 | 1621 | 0 | 0 | 0 |

**Table S10**. Hybrid demultiplexing on all cells of dataset 9pool-CA.

|  | | Hashtag demultiplexing | | | | | | | | | | |
| --- | --- | --- | --- | --- | --- | --- | --- | --- | --- | --- | --- | --- |
|  |  | Doublet | S289 | S297 | S299 | S302 | S354 | S365 | S374 | S403 | S409 | Negative |
| SNP demultiplexing | doublet | 488 | 161 | 102 | 96 | 58 | 104 | 147 | 142 | 222 | 169 | 38 |
|  | singlet0 | 134 | 39 | 27 | 34 | 26 | 27 | 42 | 35 | 63 | 58 | 7 |
|  | singlet1 | 624 | 195 | 139 | 137 | 78 | 148 | 171 | 200 | 321 | 237 | 60 |
|  | singlet2 | 371 | 178 | 62 | 83 | 44 | 60 | 106 | 123 | 164 | 142 | 19 |
|  | singlet3 | 261 | 81 | 86 | 72 | 28 | 64 | 75 | 80 | 146 | 93 | 12 |
|  | singlet4 | 462 | 129 | 108 | 103 | 60 | 87 | 120 | 146 | 195 | 225 | 34 |
|  | singlet5 | 237 | 83 | 55 | 68 | 29 | 61 | 76 | 63 | 114 | 102 | 13 |
|  | singlet6 | 372 | 106 | 91 | 82 | 41 | 82 | 90 | 152 | 178 | 126 | 20 |
|  | singlet7 | 293 | 98 | 73 | 68 | 34 | 63 | 115 | 94 | 141 | 128 | 22 |
|  | singlet8 | 570 | 216 | 141 | 105 | 53 | 105 | 138 | 158 | 355 | 224 | 41 |
|  | unassigned | 98 | 25 | 29 | 16 | 15 | 17 | 23 | 27 | 47 | 34 | 11 |

**Table S11**. Hybrid demultiplexing on dataset 9pool-CA after remove all B&T doublets.

|  | | Hashtag demultiplexing | | | | | | | | | | |
| --- | --- | --- | --- | --- | --- | --- | --- | --- | --- | --- | --- | --- |
|  |  | Doublet | S289 | S297 | S299 | S302 | S354 | S365 | S374 | S403 | S409 | Negative |
| SNP demultiplexing | doublet | 344 | 50 | 48 | 39 | 18 | 56 | 50 | 47 | 70 | 49 | 6 |
|  | singlet0 | 214 | 116 | 55 | 54 | 50 | 41 | 60 | 72 | 96 | 930 | 10 |
|  | singlet1 | 185 | 521 | 99 | 74 | 45 | 48 | 80 | 105 | 131 | 88 | 14 |
|  | singlet2 | 94 | 74 | 25 | 34 | 145 | 22 | 35 | 42 | 43 | 50 | 4 |
|  | singlet3 | 137 | 83 | 388 | 45 | 32 | 30 | 67 | 81 | 60 | 57 | 4 |
|  | singlet4 | 80 | 44 | 23 | 31 | 16 | 381 | 38 | 32 | 37 | 20 | 6 |
|  | singlet5 | 123 | 51 | 39 | 406 | 32 | 36 | 69 | 57 | 47 | 49 | 1 |
|  | singlet6 | 143 | 67 | 63 | 49 | 35 | 52 | 510 | 85 | 66 | 71 | 9 |
|  | singlet7 | 258 | 127 | 69 | 57 | 50 | 57 | 62 | 85 | 1262 | 94 | 15 |
|  | singlet8 | 168 | 94 | 61 | 63 | 31 | 57 | 90 | 573 | 89 | 77 | 11 |
|  | unassigned | 92 | 24 | 25 | 3 | 8 | 31 | 23 | 32 | 30 | 39 | 5 |

**Table S12**. Hybrid demultiplexing on dataset 9pool-CA after remove all doublets identified by cell hashing method.

|  | | Hashtag demultiplexing | | | | | | | | | | |
| --- | --- | --- | --- | --- | --- | --- | --- | --- | --- | --- | --- | --- |
|  |  | Doublet | S289 | S297 | S299 | S302 | S354 | S365 | S374 | S403 | S409 | Negative |
| SNP demultiplexing | doublet | 0 | 15 | 12 | 17 | 8 | 13 | 17 | 13 | 28 | 14 | 26 |
|  | singlet0 | 0 | 0 | 0 | 32 | 414 | 0 | 0 | 0 | 0 | 0 | 4 |
|  | singlet1 | 0 | 0 | 0 | 0 | 0 | 47 | 999 | 0 | 0 | 1 | 6 |
|  | singlet2 | 0 | 0 | 0 | 0 | 0 | 0 | 0 | 1 | 91 | 1382 | 26 |
|  | singlet3 | 0 | 0 | 23 | 809 | 0 | 0 | 0 | 2 | 0 | 0 | 7 |
|  | singlet4 | 0 | 0 | 0 | 0 | 0 | 1 | 59 | 1100 | 0 | 0 | 6 |
|  | singlet5 | 0 | 14 | 852 | 0 | 0 | 0 | 0 | 0 | 0 | 0 | 5 |
|  | singlet6 | 0 | 0 | 0 | 0 | 37 | 730 | 0 | 0 | 1 | 0 | 10 |
|  | singlet7 | 0 | 1205 | 0 | 0 | 0 | 0 | 0 | 0 | 0 | 0 | 9 |
|  | singlet8 | 0 | 0 | 0 | 0 | 0 | 0 | 1 | 71 | 1798 | 0 | 30 |
|  | unassigned | 0 | 77 | 26 | 6 | 7 | 27 | 27 | 33 | 28 | 141 | 148 |
